## Supplementary Text for "Is Vaccination a Viable Method to Control Johne’s Disease Caused by *Mycobacterium avium* subsp. *paratuberculosis*? Data from 12 Million Ovine Vaccinations and 7.6 Million Carcass Examinations in New South Wales, Australia from 1999-2009"

**S1: Introduction**

The importance of vaccination in controlling OJD was demonstrated more than 50 years ago in Iceland, where widespread clinical cases emerged in a naïve sheep population, five years after introduction of infected sheep from Germany in 1933. As they were unable to culture the strain involved, they used two bovine strains in the locally developed oil-adjuvant vaccine. Trialled in 1950 and made compulsory in endemic areas in 1966, it was reported to have virtually eliminated clinical disease. Prior to the vaccination program the average annual mortality rate from MAP in adult sheep was 8-9% and approached 40% on some individual farms [1, 2]. The organism involved was subsequently identified as an “S” strain in 2001[2, 3] .

OJD has been found at varying prevalence levels in the major sheep producing countries (New Zealand, Spain, South Africa and Europe), however it has generally been accepted as a low-grade endemic disease with a relatively low mortality rate, not warranting a coordinated government/industry control program [2].

It is considered likely that OJD had been introduced into the Central Tablelands region of the NSW HPA with imported British Breed carpet wool sheep from New Zealand in the early 1970s [2, 4].

**S2: Vaccine regulation**

Prior to January 2000 the use of vaccine was prohibited by national veterinary authorities in Australia due to lack of international research data on vaccine efficacy and concerns that vaccination would encourage a false sense of security in the minds of producers, mask the presence of “carrier” animals and interfere with the use of blood tests for diagnostic purposes. There was also concern about producer accidental self-inoculation and the presence of vaccination abscesses at slaughter. Despite these issues, the severe impact of clinical disease in many merino flocks in the NSW HPA led to “special use” approval for “Gudair®” vaccine in 2000, while live attenuated vaccines remained banned.

### The imported vaccine was also in short supply due to the long lead time in manufacturing and safety testing individual batches, as well as uncertainty about the future demand from producers.

### Vaccination was initially permitted from January 2000 on up to 50 individual properties [5]. Eligible properties were required to enrol in an “Extended Use Vaccination Trial” under a “special use permit” (Permit 3257 23 December 1999) issued by the National Registration Authority for Agricultural and Veterinary Chemicals (NRA) [6]. Eligible flocks were required to have been quarantined due to the presence of OJD, demonstrate unacceptable losses attributable to OJD (≥5% annual mortality), and be subject to a wide range of other stringent conditions. These included approval of individual properties by the NSW Chief Veterinary Officer (subject to endorsement by the Australian National Animal Health Committee), producer attendance at an OJD Vaccination Workshop and implementation of an on-farm PDMP [5, 7].

### Subsequently this “special use permit” was extended (permit 4730 4 July 2001) to include up to 150 properties in NSW as OJD continued to spread within and between flocks. Prior to November 2001, vaccinated sheep were only permitted to be sold direct to an abattoir for slaughter. Sales were subsequently allowed to other approved vaccinating properties. The vaccine was registered by the NRA for wider use in the control of OJD nationally (Permit 5489/53839) on 16 April 2002 after results of the vaccine trial [8] became available in December 2001. However it remained heavily regulated and was restricted primarily to infected flocks in the HPA[9]. In April 2003 vaccination was extended to infected flocks in NSW outside the HPA, but this remained subject to NSW Chief Veterinary Officer approval. Unrestricted vaccine access (ie. use in any flock in NSW) was approved by National Animal Health Committee in July 2003.

### Producer access to vaccine was restricted to sale through RLPBs from 2000 to December 2003 and subject to recording of both sales and property-of-use details including class and age of sheep vaccinated. The price of vaccine purchased through RLPBs was AU$1.65 per sheep from 2000-2005 and $1.85 in 2007, with the cost of administration (mustering and labor) estimated at AU$1.00. Vaccine sales by approved veterinary practices were permitted from January 2004. In August 2007 rural merchandisers were also approved to sell vaccine. For all subsequent vaccine sales, the property-of-use details were required to be documented on an internet database (Gudair® Vaccine Portal) established by Pfizer Animal Health.

**S3: Vaccine procedures**

To minimize the risk of vaccine abscesses in valuable cuts of meat [10] and the risk of self-inoculation by the vaccinator [11, 12], it was required to be administered subcutaneously in the upper neck behind the ear.

Additional safety measures included the development by Pfizer of the “Sekurus®” automatic syringe with a shrouded needle, which was only exposed when the shroud was pushed against the skin of the sheep.

To optimize the advantage of vaccination before significant exposure to MAP, it was recommended in the NRA permit as a single dose in lambs 4-12 weeks of age, however this was modified in NSW to 4-16 weeks to more closely align with age at lamb marking (tail-docking, castration and routine vaccination).

Lambs vaccinated at 4-16 weeks, or older sheep considered not to have been exposed to MAP prior to vaccination, were considered low risk ‘approved vaccinates” and were subject to less strict trading regulations.

**S4: Property disease management plans (PDMPs), financial assistance and producer workshops**

The concept of formal PDMPs was promoted through a series of workshops commencing in February 2002 (581 producer attendees) with vaccination as a key control measure, complemented by a range of other management strategies based on an evolving understanding of the epidemiology of OJD [7, 13].

Financial assistance was provided to implement PDMPs (funding various strategies, primarily vaccination) over a 3-year period following PDMP approval. Sourced from a NSW sheep industry levy, it was made available from May 2002 at AU$5 per head based on the number of sheep in the flock (up to a maximum of 5000). This was only accessible to producers with flocks confirmed as OJD infected prior to 4 November 2002, who attended workshops (179 producers) and developed a formal PDMP. The great majority of this funding was utilized for purchase of vaccine in 2002 and 2003. A further series of 14 workshops (454 attendees) was held within the HPA and MPA between July and August 2003 to promote better management and control of OJD, with the emphasis on vaccination [13, 14].

Issues promoted extensively to livestock producers throughout these workshops included:

1. The importance of prevalence areas based on abattoir monitoring in assessing regional risk,

2. The results of the vaccine trial with 90% reduction in clinical disease and reduced excretion of MAP [8],

3. The importance of safe vaccination technique, and

4. On-farm management tools based on the emerging understanding of the epidemiology of OJD [15]. These strategies included enhanced biosecurity, selective culling of high-risk groups, segregation of test negative groups, partial de-stocking, artificial breeding, water and land management, protection of young susceptible lambs, spelling of land to reduce contamination levels.

These workshops, combined with the reporting to producers of positive consignments detected by abattoir monitoring (including the percentage of animals with lesions attributable to OJD), led to a marked upsurge in the use of vaccine by sheep producers.

### S5: Inspection, sampling & data collection

Consignments were usually slaughtered as a single contiguous line of sheep passing down the slaughter chain. Data from each consignment monitored were recorded by the inspectors, entered in the central database and reported individually to producers. However, to facilitate processing, larger consignments of sheep were often separated by abattoir management into smaller “*split*” consignments and killed separately, sometimes on different days. These “*split”* consignments were identified by the inspectors based on a common Lot No., Owner and Locality/PIC, and were inspected and recorded in the database as separate but “linked” consignments. In order to minimize laboratory costs, a maximum of three samples were usually collected from the combined “split” consignments. Positive histopathology results in any “split” consignment were applied across all “linked” consignments, irrespective of the presence or absence of gross lesions.

The total number of sheep with lesions truly attributable to OJD in “Positive” consignments was estimated by *pro-rata* correction (based on the results of histopathology) of the number of animals with lesions grossly resembling OJD as originally reported by the inspector.

For example, 300 sheep inspected, 10 lesions detected, 3 sampled for histopathology:

- 3/3 POS – **“corrected”** lesions = 10
- 2/3 POS – **“corrected”** lesions = 7
- 1/3 POS – **“corrected”** lesions = 3
- 0/3 POS – **“corrected”** lesions = 0, consignment is OJD “Negative”

**S6: Feedback to producers**

Prior to introduction of the NLIS, verification of the identity of the owner of each histopathology “Positive” consignment was based on a detailed audit of abattoir consignment records and required labour-intensive retrospective visits to the abattoir. Due to the impact of quarantine on properties detected infected, there was zero tolerance for error in confirming the owner of the sheep monitored.

The owner verification and notification process for positive consignments was streamlined with the progressive implementation of NLIS in Australia. This included the introduction of the compulsory National Vendor Declaration forms which accompanied the sheep to the abattoir, allowing recording of the PIC for all consignments at the time of slaughter.

**S7: Discussion**

**S7a -** **Importance for the Merino wool and sheepmeat industry**

In 2009 it was estimated that there were 77 million sheep in Australia with 34% (26.1 million) in NSW. These comprised about 84.3% (64.8m) Merino, 11% (8.5m) 1^st^ Cross Merino/British Breed and 4.8% (3.7m) Crossbred/British Breed sheep. The respective numbers for NSW were 84%, 11.7% and 4.4% [16].

**S7b - Economic and social impact of OJD**

Prior to the availability of vaccine, the psychological impact on producers forced to watch increasing numbers of animals slowly waste away and die was heart wrenching in the absence of effective control and treatment measures. Owners described affected animals initially lagging behind other sheep in the mob when moved, then progressively deteriorating in condition over a period of 2-3 months from onset of first signs. One owner described the distress each morning of finding crows perched on fence posts waiting for the next sheep to die.

Initial losses in a flock tended to be in purchased infected sheep or in older homebred sheep exposed to low levels of contamination years earlier. However, once OJD was well established in a flock (losses >5% annually) the increasing level of contamination and exposure of lambs at an early age led to clinical disease in progressively younger adult age groups. Progression of disease is dependent on dose, period of exposure and age when exposed. While sheep can become infected at any age, young lambs are more susceptible than hoggets with adults relatively more resistant [15], a similar pattern of age resistance as that to internal parasites. Late- stage clinical cases excrete ~10^8^organisms/gm of feces [8] resulting in high levels of pasture contamination once clinical disease commences. Many heavily infected flocks suffered severe losses in maiden ewes (3 years old after lambing). This rendered it difficult for those flocks to remain self–replacing due to loss of flock structure and genetics [17]. Failure to recognise the importance of increasing resistance with age contributed to accelerating animal-level prevalence when young lambs grazed pasture recently grazed by infected adults.

The economic impact of quarantine on **sheep studs** suspected or known to be infected with OJD was particularly devastating. Their reputation was destroyed, their high-value trading ceased and they were forced into less profitable commercial wool and meat production [5]. Commercial flocks in quarantine were also severely affected, with reduction in land values by banks having a major impact on their viability. This led to increasing levels of producer resistance and regulatory non-compliance in affected regions.

District Veterinarians played the major role in liaising with producers whose flocks were impacted by OJD, including notifying producers of OJD positive abattoir monitoring results. Prior to the approval to use Gudair®, the NSW regulatory program was perceived by many producers to be aimed at eradication. This meant that roll-out of the vaccination program in the HPA was complicated by negative attitudes among many quarantined infected flocks that had suffered program-induced economic loss. This made vaccination uptake all the more remarkable and is testimony to the dramatic clinical improvement seen in many heavily infected flocks. The impact of PDMP’s on prevalence was also considered minor overall by many District Veterinarians. Most producers routinely vaccinated at lamb-marking or weaning without significantly modifying management procedures. A far more likely adjunct to the positive impact of vaccination on prevalence was the incorporation of crossbreeding programs, involving older Merino ewes with British Breed rams, in many commercial HPA flocks. This move away from Merino wool production was a response to increased sheepmeat prices. This meant that at least a proportion (generally under 20%) of their infected lambs were slaughtered prior to any potential ongoing transmission, rather than being retained for wool production (J. Evers, pers. comm.).

**S7c – Factors in disease spread**

Regulatory movement controls, plus increasing publicity on the regional distribution of OJD (including declaration of prevalence areas) undoubtedly played a major role in slowing down the inter-regional spread of OJD. Concurrently, there was a strong financial imperative driving the trend towards ultrafine wool production in the Australian sheep industry beginning in the 1990s. This resulted in movements of sheep to new regions from studs or elite commercial flocks, many from areas subsequently identified as High Prevalence. This increased the risk of unintended spread if the stud or flock of origin was sub-clinically infected with OJD. Producers came to realise that the risk of introducing infection was directly related to the numbers of sheep purchased and the prevalence within the flock of origin [17].

Many studs joined the national OJD Market Assurance Program [5] with testing (later complemented by vaccination) and attention to enhanced bio-security providing assurance that they were a source of low-risk sheep. However, flocks undergoing assurance testing were quarantined if found infected, and this proved a major disincentive to testing (including avoiding abattoir monitoring) for many stud producers. The introduction of prevalence areas severely restricted the trading of untested flocks from the HPA to lower risk areas.

**S7d - Cost of vaccination**

While only lambs to be retained on farm (as replacement ewes or castrated wethers for wool production) were targeted for vaccination, the up-front cost of vaccination was significant. It represented ~2.5% to 5% of the value of a 1.5 year-old re-stocker merino ewe, typically selling for AU$50 to AU$100 (average prices between 1999 and 2009). However, as the vaccine is a single dose for life, the average annual cost to vaccinate a ewe lamb, subsequently culled at 5 years of age, was AU$0.50 to $0.60, i.e. ~0.5% to 1% of the value of a replacement ewe per year.

**S7e - Vaccinating infected sheep – concerns regarding exacerbation of disease**

There was initial concern from national veterinary authorities that vaccination would exacerbate the progression of clinical disease leading to a rise in mortalities. Hence there was a reluctance to register the vaccine for widespread use until it had been fully clinically evaluated. While no problems were experienced following vaccination of lambs < 14 weeks of age in heavily infected flocks commencing January 2000, serious concerns remained about vaccinating older sheep. Fortunately, this question was answered by late- 2000 following completion of the initial vaccination of adult sheep in a “Whole-of-Flock” vaccination trial reported by Windsor (2005) [18]. The 10,000 head fine-wool Merino flock in the HPA was suffering extreme losses attributable to OJD – the owner estimated [18, 19] annual mortality of 25% in 1999 compared with 3-5% before confirmation of OJD on the property in 1996. This extreme mortality rate, from a long incubation and generally progressive disease, is consistent with a very high proportion of the flock being heavily infected at the time of initial vaccination between May and August 2000.

In combination with major changes in management, to minimise pasture contamination and decrease exposure of more susceptible age groups (including elimination of cell-grazing with very heavy stocking rates), the estimated annual mortality rate attributable to OJD declined from 19.0% in year 0 (August 2000-August 2001) and 18.6% in year 1 to 12.2% in year 2 and 1.4% in year 3 [18]. (**Definition: *Cell Grazing*** *– Mobs of up to 5000 sheep were grazed on 10Ha paddocks with a change of paddocks every 2-3 days*). However, Windsor [18] considered that the management changes (PDMP) were likely to be more important than vaccination in reducing mortalities dramatically by year 3. In contrast, we consider the role of vaccination, both therapeutic and preventative, as likely to have been critical in markedly reducing mortalities. This would be consistent with the findings of a 90% reduction in mortalities in the initial Gudair® vaccine trials [8], the reduction from 7.6% to 0.1% in estimated animal-level prevalence in 12 vaccinating flocks monitored over 10 years [20] and the findings in the current study.

**S7f - Vaccine – preventative or therapeutic?**

There were regular reports from OJD abattoir inspectors that sheep with advanced lesions in heavily infected consignments, that had been recently vaccinated as adults, showed mesenteric lymph nodes that appear somewhat shrunken (considered less “active” by the inspectors). This contrasts with the enlarged, oedematous lymph nodes usually seen in heavily infected consignments of unvaccinated sheep and suggests that vaccination even in the late stages of sub-clinical infection may have a therapeutic effect (W. Gilbert, pers.comm.).

**S7g - “Herd” immunity**

Many producers undertaking adult or “whole-of-flock” vaccination felt that, despite the cost, they were taking the best possible approach to rapidly maximising the level of herd/flock immunity and minimizing the longer-term risk from exposure of any uninfected sheep. This decision was often encouraged by accelerating mortality rates that threatened the financial viability of the enterprise, as well as major producer concerns about animal welfare issues.

It was psychologically distressing for producers forced to watch a continual “dribble” of individual sheep waste away and die over a period of weeks or months, without any effective treatment or alternative prevention mechanisms. Many producers cited “peace of mind” as an important contributing factor for pursuing “whole-of-flock” vaccination – confident that “they had done all they could” to control OJD in their flock [21]. In contrast, prior to the advent of abattoir monitoring, producers with a low level of sub-clinical disease in their flock, and no evidence of clinical disease, found it very difficult to accept that their flock was infected, and were hence reluctant to embrace vaccination.

Further research on whole-of-flock vaccination in the face of heavy challenge is required to test the disease control, therapeutic and financial benefits of this strategy.

Unfortunately, due to logistical constraints in the abattoir, and despite mandatory identification of vaccinated sheep by ear-tag, we were unable to reliably classify the vaccination status of all individual consignments (or sheep), many of which included a mix of vaccinated and unvaccinated animals.

**S7h - Sensitivity and specificity**

A research trial was incorporated into the routine abattoir monitoring in 1999-2000, to determine the sensitivity and specificity of visual inspection with follow-up histopathology on 3 animals [22].

Inspectors were blind to the OJD status of properties with consignments included in the trial, which were examined over a 6-month period. Gross lesions were detected in 34/35 (97%) consignments from selected known infected properties. Microscopic lesions diagnostic for OJD were identified in 31 (91%) of the 34 consignments detected by inspectors while the remaining 3 (9%) yielded inconclusive histopathology. The average consignment size was 330 sheep (range 50-683). The average proportion of sheep in these 34 consignments with gross lesions suggestive of OJD, as reported by inspectors, was 21% (range <1% to >90%). No visible lesions of OJD were detected in 7 of the 9 negative controls (from properties not known to be infected with OJD). In the remaining 2 controls, however, inspectors reported 2% and 4% of sheep with suspect lesions, but all fixed tissue samples submitted were negative on histopathology.

The estimated consignment sensitivity from this data was 97% (95% confidence intervals: 91.5% - 100%). They suggested a practical working sensitivity estimate of 90% for visual/tactile examination of viscera for lesions suggestive of OJD in consignments of >300 adult sheep. This supported the high sensitivity of abattoir surveillance, as a screening test for the selection of samples prior to a definitive histopathology test, for any sheep population in which OJD has been recognised for many years. The limited data for “low prevalence flocks” (<2% within flock prevalence) obtained in that trial suggested that visual/tactile examination of viscera may be a more sensitive technique for detection of low prevalence infected consignments than was previously believed, and was probably more sensitive than the estimate of 30% at that time. They recommended that further investigation of the utility of this method of abattoir surveillance for OJD in low prevalence flocks and low prevalence areas was warranted [22].

The performance of individual inspectors in the 11 NSW abattoirs in the present study was analysed for specificity. A total of 36 trained inspectors participated in the 10-year study located variously in abattoirs predominantly servicing either the HPA, the LPA or a mix of prevalence areas. A preliminary report [23] confirmed that 19 of the inspectors monitored 98% (5,658/5,792) of consignments sampled (range 22-810) **sourced from all of NSW** with an average inspection rate of 75% (range 54-95%) and a **relative consignment specificity of 79%** (range 52-99% - depending on the predominant prevalence area monitored). Similarly, these 19 inspectors submitted 12,303/13,101 (94%) samples with a **relative sample specificity** **of 76%** (range 43%-97%). In contrast, the 17 minor inspectors had relative consignment and sample specificities of 85% and 85% respectively (they were predominantly monitoring sheep from the HPA). The relative specificities were reduced by deliberate oversampling of animals with any suspicious gross pathology in consignments from the LPA, aimed at maximising sensitivity while limiting samples to three viscera.

The equivalent relative specificities for those inspectors examining consignments **sourced from the HPA,** where oversampling was limited, were **consignment specificity of 86%** (range 67%-100%) and **sample specificity of 82%** (range 55%-99%).

**S7i - Saleyard consignments**

To ensure more comprehensive monitoring in the LPA where many producers routinely sell through local or regional saleyards, mixed saleyard consignments were routinely monitored. When positive, contributing flocks were identified (PIC, NLIS) and a detailed risk assessment undertaken, followed by on-farm testing where justified [24].

In the latter stages of this study (2007-2008) export abattoirs introduced individual animal identification based on the ear-tag NLIS PIC recorded at the time of slaughter. The PIC for each animal was displayed on a monitor adjacent to the viscera inspection site, enabling the inspector to record the PIC for individual animals with lesions in mixed saleyard consignments as well as to eliminate the risk of mis-identification of sheep in direct consignments. The results from mixed saleyard consignments have not been included in the current study.

Some studs and other commercial producers deliberately avoided monitoring by consigning sheep through saleyards, or to smaller abattoirs not currently being monitored, to avoid the impact of regulatory controls on their enterprise if detected infected. The introduction in 2008 of abattoir assurance credits to facilitate improved trading opportunities for flocks that monitored negative, encouraged many reluctant producers to embrace abattoir monitoring. Flocks were eligible for Abattoir 150 or Abattoir 500 status if 150 or 500 sheep were monitored negative over 12 months or 2 years respectively. It provided further support for the Assurance Based Credit (ABC) producer driven risk-based trading system initially introduced across Australia in 2003 [14, 25].

**S7j - Vaccine abscesses**

Monitoring direct consignments of adult sheep from the NSW HPA between 2006 and 2012 for vaccine abscesses confirmed a high level of compliance, with 1.9% (101/5,212) of consignments affected, involving 2.6% of sheep with abscesses (33,431/1,282,065). They were derived from 4.1% of properties monitored (79/1934 PICs) with the great majority of individual properties detected only once (63 properties), and a small number of properties with multiple detections - twice (11), 3 times (4) or 4 times (1) (I.Links, unpublished observations). The majority of these animals would have been vaccinated during the period of this study (2002 – 2009). Vaccine abscesses were generally attributable to faulty operator procedure resulting in intramuscular rather than sub-cutaneous injection, with the oil emulsion vaccine trapped by the fascia. Nonetheless, producers (and the sheep industry) need to be proactive in ensuring that faulty vaccination technique is not leading to development of vaccine abscesses, particularly where vaccination is being undertaken on a property for the first time. It is feasible that some vaccination techniques are unsatisfactory, possibly introduced due to a desire to protect the operator from accidental self-inoculation.

**S7k - Potential role of vaccination in controlling BJD in cattle**

In an endeavour to overcome some of the perceived constraints of killed oil emulsion MAP vaccines (such as vaccination abscesses, risk of accidental self-inoculation and interference with diagnosis of tuberculosis in dairy cattle), recent trials in the USA [26, 27] have evaluated five live attenuated potential vaccine strains (administered orally) against a commercial heat killed oil adjuvant vaccine injected subcutaneously (Silirum®), using a standard goat challenge model. Only the control vaccine (Silirum^®^) showed a clear reduction in lesion score, MAP faecal shedding and tissue colonization. Their results suggest that although vaccination with Silirum^®^ does not prevent infection or eliminate MAP faecal shedding in goats, it reduces presence of JD gross and microscopic lesions, and slows progression of disease. These findings are consistent with those previously reported in trials with Gudair® in sheep in Australia [8] and in the present study.

While the superior performance of killed vaccine may possibly be attributed to the route of administration, it is also likely that long term, possibly lifelong, persistence of the oil adjuvant vaccine continually primes the cellular immune system. MAP are reported to survive and multiply after invasion of macrophages by preventing phagosome maturation. This “immune subversion” is an active process which prevents initial killing of MAP by switching from an appropriate Th1-like T-cell response to an ineffective Th2-like response, enabling the MAP organisms to safely multiply intra-cellularly and hide from the immune system. The cycle of infection continues following liberation of MAP from apoptotic macrophages and subsequent sequestration in new macrophages – a short period during which they are once again exposed to the immune system [28]. If such is the case, it would support the concept that vaccination of animals incubating MAP with a long-term persistent vaccine could be therapeutic, potentially tipping the balance in favour of the host immune system rather than the microbe. It could thus delay or even reverse, the progression of infection to microscopic and gross pathological changes and ultimately clinical disease.

As in the present study, vaccination of older age groups in heavily infected cattle herds would be expected to enhance control of infection by increasing herd immunity more rapidly than would occur if vaccination was restricted solely to young calves.
